# Benchmarking nanopore-based strategies for antimicrobial resistance prediction

**DOI:** 10.64898/2026.04.06.716670

**Authors:** Natalie Ring, Alison S. Low, Rhodri Evans, Marianne Keith, Gavin K. Paterson, David Gally, Tim Nuttall, Dylan Neil Clements, J. Ross Fitzgerald

**Author notes:** **Corresponding author and email address**, Natalie Ring. **Repositories:** Sequencing data for all samples sequenced for this study are available at the NCBI under BioProject ID PRJNA1292816 (SRA accessions for all datasets used here are available in Supplementary Table S1).

## Abstract

2.

Antimicrobial resistance (AMR) presents a pressing need to ensure that the right antimicrobials are used to target the right microbes at the right time. Ideally, the appropriate antimicrobial is selected after patient samples have been cultured and assessed with antimicrobial sensitivity testing (AST). However, the time needed for culture-based diagnosis leads to immediate empirical treatment, often with broad-spectrum and/or high-tier antimicrobials. Direct nanopore metagenomic whole genome sequencing to identify pathogens and predict their antimicrobial resistance is a rapid and patient-side alternative. A limitation of this approach is potential inconsistencies in *in silico* predicted AMR phenotypes. Here, we benchmarked the current performance of *in silico* AMR prediction strategies for nanopore-generated long read data. Using nanopore data paired with AST phenotyping for 201 samples representing 27 bacterial species, we assessed the impact of basecalling mode, data volume, and assembly strategy, and compared the performance of eight *in silico* AMR prediction tools with seven AMR databases. We found that basecalling accuracy mode does not significantly affect the overall accuracy of *in silico* AMR predictions, but assembly strategy and data volume both do. Prediction tools using the ResFinder database scored best for balanced accuracy (0.80 ± 0.02 for both ResFinder and ABRicate), whilst DeepARG scored best for sensitivity (0.65 ± 0.03); predictions were more accurate for some antibiotic classes and genera than others. However, even the best performing *in silico* AMR prediction strategy missed some resistance identified by lab-based AST. We conclude therefore that, currently, *in silico* AMR prediction can supplement lab-based AST, but cannot yet replace it.

**Impact statement:** Antimicrobial resistance (AMR) is threatening modern standards of human and veterinary healthcare. Rapid and patient-side diagnostic tests are needed to diagnose bacterial infections and allow clinicians to select effective antibiotics. Current tests based on bacterial cultures take several days, which may delay diagnosis and treatment, or lead to inappropriate “just in case” treatment while waiting for the results. In contrast, nanopore metagenomic whole genome sequencing can identify bacterial infections and predict which antibiotics will be effective in minutes to hours. However, the accuracy of these tests is uncertain. We therefore compared the performance of eight AMR prediction tools and seven databases of AMR determinants, using 201 bacterial samples with known antibiotic susceptibility and/or resistance. We found that the sensitivity (i.e. false negative rate), specificity (i.e. false positive rate) and overall accuracy of the tools and databases varied. In particular, even the best performing AMR prediction methods missed some AMR. Therefore, while these tools are useful for rapid and patient-side diagnosis and treatment decisions, they still have limitations and should be used alongside bacterial cultures and antibiotic sensitivity testing.

**Data summary:** Sequencing data for the samples sequenced for this study are available at the NCBI under BioProject ID PRJNA1292816 (SRA accessions for all datasets used here are available in **Supplementary Table S1**).

All commands and code used can be found at: https://github.com/nataliering/nanopore_AMR_tools/

**The authors confirm all supporting data, code and protocols have been provided within the article or through supplementary data files.**

## 5. Introduction

Antimicrobial resistance (AMR) is an immediate and ongoing global threat that will impact health, food security and economic development (1, 2). Over eight million annual AMR-associated deaths are forecasted by the year 2050 (3). Inappropriate and indiscriminate antimicrobial use exerts a selection pressure favouring the development and dissemination of bacterial AMR (4). To avoid this, we need to use the right antimicrobial, for the right infection, at the right time. This is a one health challenge that encompasses human, production animal and companion animal healthcare.

Faster diagnosis to confirm a bacterial infection and determine its antimicrobial susceptibility will improve antimicrobial stewardship. The current gold-standard methods involve culturing samples to identify the bacteria, followed by antibiotic sensitivity tests (ASTs). Commonly used ASTs include disk diffusion, minimum inhibitory concentration (MIC) test strips and broth microdilution assays (5, 6). These tests require a minimum of 16 hours, but it often takes significantly longer to obtain the results. Immediate empirical antimicrobial treatment may be unnecessary (where there is no bacterial infection) and/or may employ unnecessarily broad-spectrum or high-tier antimicrobials through fear of compromising patient outcomes. Excessive and inappropriate antimicrobial use is a major driver of AMR (4). In addition, culture-based ASTs are effectively “a sample of a sample” (i.e. patient sample – culture – culture sample – AST) and favour less fastidious bacteria. AST results may not therefore reflect the whole bacterial population and antimicrobial susceptibility patterns in the patient’s infection.

Metagenomic whole genome sequencing (mWGS) using Oxford Nanopore Technologies’ (ONT) long-read sequencers is a rapid culture-free technique that involves extraction and sequencing of all genomic DNA (gDNA) in a sample. The unbiased DNA sequencing data can be used to identify pathogens and to detect genes and mutations associated with AMR in a little as 10 minutes (7, 8). Progress on developing mWGS-based rapid diagnostics pipelines has accelerated in recent years, and ONT’s platforms have been used to pilot rapid diagnostic methods for mWGS from clinical samples such as sputum, respiratory fluid, blood, urine, and skin swabs (9–24).

Robust, reliable and reproducible bioinformatic tools and pipelines will be vital to implement rapid mWGS-based diagnostics in clinical settings (25). In 2020, a multi-laboratory study found wide discordance between the AMR phenotypes predicted by different laboratories using the same genomic datasets (26). This discordance occurred as a direct result of differences in how each laboratory chose to process the data, and the bioinformatic tools and databases used for the AMR predictions. Similarly, a comparison of tools to identify AMR from Illumina short-read datasets showed wide variability in their sensitivity, specificity and balanced accuracy (27). No single tool performed equally well across every antibiotic class; some performed very well in terms of sensitivity but relatively poorly in terms of specificity.

Here, we analysed the performance of eight different AMR detection tools: ABRicate (28), StarAMR (29), ResFinder (30), AMRFinderPlus (31), c-SSTAR (32), RGI (33), DeepARG (34) and AMR++ (35) on datasets produced using the ONT sequencers that would be used for mWGS-based rapid diagnostics. We sourced a collection of 201 ONT datasets with matched lab-based AST phenotypic AMR profiles, representing 27 different bacterial species. We included tools which require the assembly and annotation of sequencing reads into consensus contigs as well as read-based tools. The tools use different methods to detect AMR determinants, including *k*-mer-based alignment, Hidden Markov Models (HMM), and deep machine learning. Some tools offer a choice of AMR databases (e.g. ABRicate) whilst others are based around their own database (e.g. ResFinder, DeepARG). We tested all possible combinations and options for the tools, as well as addressing issues such as the impact of data volume, the downstream effects of different basecalling accuracy options, and the optimal assembly strategy. We also compared the performance of tools at the level of antibiotic class and individual antibiotics.

## 6. Methods

### Data collection and curation

An NCBI PubMed database search was conducted in May 2025, using various combinations of the terms “nanopore”, “AMR”, “MinION” and “AST”. The resulting literature hits were manually reviewed to determine if 1) their raw sequencing data was publicly available and 2) their datasets included *vitro* antimicrobial sensitivity testing (AST) results. If a paper fulfilled both of the above criteria, the relevant datasets were downloaded, and the following metadata was recorded for each sample (full data in **Supplementary Table S1**): sample accession, sample type, MinION flow cell version, library preparation kit, basecaller used (and, where available, version and accuracy mode), species identified in original publication, AST results, and original publication reference.

### Comparing monocultured isolates with metagenomic samples

The datasets gathered here were separated into two main groups: monocultured isolates, and metagenomic samples with potential host contamination or containing multiple species. The monoculture group (151 samples) was used for the majority of the analysis described below, including comparison of basecalling and assembly strategies, investigation into data volume, comparison of tool and database performance, comparison of prediction accuracy across different antibiotic classes, and comparison of prediction accuracy across different genera. The metagenomic group (50 samples) was used to assess whether the tool performance patterns identified by the monoculture analysis were also observed for the more “difficult” sample types.

### Read preparation

Data were downloaded in fastq format from the NCBI sequence read archive (SRA) using fasterq-dump v2.11.3 (36). NanoStat v1.6.0 (37) was used to determine mean read length, number of reads, and overall number of bases. Datasets were excluded if their mean read length was less than the standard nanopore quality cut-off of 1,000 bp (38–41) or their overall number of bases was less than 25 Mb (around 5**×** coverage of the “average” monocultured bacterial genome (42–45)). A set of 64 additional isolates (**Supplementary Table S2**) were sequenced specifically for this study, representing species not otherwise found in the above literature search. See **Supplementary Materials SM1** for full DNA extraction, library preparation and sequencing methods, and **Supplementary Table S1** for sample accessions.

Porechop v0.2.4 (46) was used to trim adapters from the fastq read sets which passed quality control (QC), and Filtlong v0.2.0 (38) was used to filter longer read sets down to 0.5 Gb of the longest and highest quality fastq reads. Where an AMR tool based its predictions on raw fastq reads alone, the output Filtlong reads were used.

### Genome assembly and draft annotation

The trimmed reads were assembled with two different long-read assembly tools, Minimap2 v2.24-r1122/Miniasm v0.3-r179 (47, 48) and Flye v2.9 (49). The Flye assembly was polished using Medaka v2.0.1 (50), and the Flye, Miniasm and Medaka assemblies were all annotated with Prokka v1.14.6 (51). All three assemblies (Miniasm, Flye and Medaka), along with the raw reads and annotations where applicable, were used to assess the performance of the tested AMR tools to determine the optimal assembly strategy. More details about each assembly tool can be found in the **Supplementary Materials** (**SM2**).

### AMR tools and databases tested

We tested a variety of assembly- and read-based tools, with every relevant database available to each tool. The tools and databases are summarised in **Table 1** and more details are in the **Supplementary Materials** (**SM3**). Commands used for each tool are available from: https://github.com/nataliering/nanopore_AMR_tools/.

**Table 1.** Antimicrobial resistance detection tools and databases tested here.

| Tool | Version | Resistance database(s) | Resistance database version | Input file(s) |
| --- | --- | --- | --- | --- |
| ABRicate (28) | 1.0.1 | NCBI(31),<br>CARD(33),<br>ResFinder(30),<br>ARG-ANNOT(53),<br>MEGARes<br>2.0(54) | 14 Jan 2025 | Assembly |
| AMRFinderPlus (31) | 4.0.23 | NCBI | 2025-03-25.1 | Assembly and annotations |
| ResFinder (30) | 4.7.2 | ResFinder &<br>PointFinder | 2.6.0 &<br>4.1.1 | Reads or assembly |
| c-SSTAR (32) | 2.1.0 | ResGANNOT | NA | Assembly |
| RGI (33) | 6.0.4 | CARD | 4.0.1 | Assembly, or assembly and annotations |
| StarAMR (29) | 0.11.0 | ResFinder | 2024-08-06 | Assembly |
| DeepARG (34) | 1.0.2 | DeepARG DB | NA | Reads, or assembly and annotations |
| AMR++ (35) | 3.0.6 | MEGARes | NA | Reads |

Abbreviations: AMR, antimicrobial resistance; NCBI, National Center for Biotechnology Information; CARD, Comprehensive Antibiotic Resistance Database; ARG-ANNOT, Antibiotic Resistance Gene-ANNOTation; RGI, Resistance Gene Identifier; DB, database; NA, not available.

For tools which used assembled contigs for AMR prediction, the Miniasm, Flye and Medaka assemblies were all tested with each tool/database combination. See **Supplementary Table S3** for a full list of the 42 tool/database combinations.

Some of the tools tested here, such as ResFinder, offer species-specific AMR prediction for a limited number of species, facilitating the detection of AMR caused by species-specific point mutations. Where a sample was known to contain one of these species, all possible combinations were also trialled with this mode. **Supplementary Table S4** shows which tools were used with which species. Metagenomic samples containing multiple species were not included in the species-specific tests.

HAMRonization v1.1.9 (52) was used to convert the results of all tools into a consistent tabular form. See **Supplementary Methods SM4** for details of the additional steps which were taken to ensure standardisation of the way results were reported across different tools.

### Calculating performance metrics

For each sample, the original AST results were collated to create a per-antibiotic “phenotypic AMR” table (**Supplementary Table S5**). In addition, the Comprehensive Antibiotic Resistance Database (CARD) ontology (33) was used to assign antibiotics in our phenotypic AMR table to their drug class: aminoglycosides, beta-lactams, fluoroquinolones, glycopeptides, macrolides-lincosamides-streptogramins (MLS), phenicols, sulfonamides, tetracyclines, trimethoprim (diaminopyrimidines), and polypeptides. Rarer antibiotic classes were excluded from analysis, due to insufficient numbers of resistant samples in our dataset. **Supplementary Table S6** shows which antibiotics were included in each class and which classes/antibiotics were excluded. A second table was then produced, which listed each sample’s phenotypic AST results in terms of drug class (**Supplementary Table S7**). For each sample in our phenotypic AMR tables, each antibiotic or class was assigned “R” (resistant), “S” (susceptible) or “NA” (not tested) based on the AST results from the original publications. Where a result in the phenotypic AMR table was “I” (Intermediate), it was considered to be resistant.

Each set of results was then analysed at the level of specific antibiotic and antibiotic class. The tool results were compared with our phenotypic AMR table, and each antibiotic/class was scored as True

Positive (TP), False Positive (FP), True Negative (TN) or False Negative (FN) for each sample. A TP occurred when both the phenotypic AMR result and the *in silico* tool prediction were “R” for an antibiotic/class for that sample. A FP occurred when the phenotypic result was “S” but the tool predicted “R”. A TN occurred when both the ground truth result and the tool prediction were “S”. A FN occurred when the phenotypic result was “R” but the tool predicted “S”. Where an antibiotic or class was not AST tested for a sample (“NA”), that antibiotic or class was excluded from the results comparison.

Performance metrics were then calculated for each tool according to the equations shown in **Table 2**.

**Table 2.** Equations used to calculate our performance metrics.

| Metric | Equation |
| --- | --- |
| Sensitivity | $\frac{TP}{TP + FN}$ |
| Specificity | $\frac{TN}{TN + FP}$ |
| Balanced accuracy | $\frac{sensitivity + specificity}{2}$ |

Abbreviations: TP, true positive; TN, true negative; FP, false positive; FN, false negative.

### Statistical analysis and data visualisation

All statistical analyses were performed in Python v3.10.14 using pandas v2.3.0 (55, 56), SciPy v1.15.3 (57) and scikit-posthocs v0.11.4 (58). Performance metrics were calculated for each comparison group as described above. Normality was assessed using the Shapiro-Wilk test (59) implemented with scipy.stats.shapiro, and homogeneity of variance was assessed using Levene’s test (60) implemented with scipy.stats.levene. Because the performance metrics were bounded between 0 and 1 and did not consistently meet parametric assumptions, non-parametric tests were used for group comparisons. Overall differences among groups were assessed using Kruskal-Wallis tests (61) implemented with scipy.stats.kruskal. Where Kruskal-Wallis tests indicated significant differences, pairwise post-hoc comparisons were performed using Dunn’s test (62) implemented with scikit_posthocs.posthoc_dunn, applying Bonferroni correction to account for multiple pairwise comparisons. Statistical significance was defined as p < 0.05.

Figures were generated in R v4.1.2 using ggplot2 v3.3.6 (63), readxl v1.4.0 (64), dplyr v1.0.9 (65), tidyr v1.2.0 (66), stringr v1.4.0 (67), forcats v1.0.0 (68) and ggpubr v0.4.0 (69). Summary tables were imported using readxl::read_excel and plotted using ggplot2. Bar plots were produced with geom_bar, error bars with geom_errorbar, and heatmaps with geom_tile. Multi-panel figures and shared legends were arranged with ggpubr::ggarrange, and Dunn’s post-hoc significance annotations were added using ggpubr::stat_pvalue_manual. Final figures were exported using ggplot2::ggsave.

### Comparing super accuracy, high accuracy and fast basecalling mode

A subset of 64 monoculture samples (all those for which raw data was available, **Supplementary Table S2**) were used to conduct a comparison of the accuracy of AMR prediction when using different basecalling accuracy modes. Where necessary, the unbasecalled fast5 files were converted into pod5 format using pod5 v0.2.15 (70). The Dorado standalone basecaller (versions 0.9.6 and 1.0.2 for data from SQK-RBK004 and SQK-RBK114.24 respectively) was then used to basecall the reads with each of the three accuracy modes: fast, high accuracy (HAC) and super accuracy (SUP), creating three basecalled read sets per sample. Each read set was processed through the data preparation, AMR prediction, and analysis pipeline as described above and performance metrics were stratified by basecalling accuracy mode.

### Assessing the effect of data volume on AMR prediction accuracy

A subset of 61 samples (all those with at least 500 Mb of starting data, **Supplementary Table S8**) were used to assess the effect of initial data volume on the eventual accuracy of AMR prediction. Each read set was processed through the data preparation, AMR prediction, and statistical analysis pipeline as above, except that instead of filtering to 500 Mb with Filtlong the read sets were filtered to 250 Mb, 100 Mb, 50 Mb and 25 Mb. Including the read set originally filtered to 500 Mb, this produced five different sets of AMR predictions, representing 100**×**, 50**×**, 20**×**, 10**×** and 5**×** coverage of the “average” length bacterial genome (42). The performance metrics for each level of filtering were compared in the same way as the basecalling modes, as described above.

### Comparing older and newer sequencing data

The performance metrics for a subset of 28 samples (**Supplementary Table S9**) were used to compare the accuracy of AMR predictions made from older and newer sequencing data. 14 samples sequenced with R10.4.1 chemistry and basecalled with Dorado were compared with 14 samples sequenced with R9.4/R9.4.1 chemistry and basecalled with Albacore or Guppy. The samples were species-matched between R10 and R9 as far as possible, according to the samples available in our dataset: 5 *Enterococcus*, 3 *Pseudomonas* and 6 *Staphylococcus* samples for each chemistry.

## 7. Results

### A curated dataset of 201 samples to evaluate AMR prediction tools

A total of 201 samples met our selection criteria of having both ONT reads and AST phenotypes available, as well as passing our QC process (12, 71–83). **Supplementary Table S1** gives the full metadata of all analysed samples and **Supplementary Table S10** gives the references of the original papers. The samples were sequenced on a variety of different flow cell types (R9.4.1, R9.5, R10.4.1), sequencing libraries were prepared using a variety of kits (for example, SQK-RPB004), and basecalling was performed with a variety of different basecallers (Albacore, Guppy, Dorado) and basecaller software versions. Starting volumes of data ranged from 28 Mb to 16.5 Gb, with a mean of 662.51 Mb, median of 175.90 Mb and standard deviation of 2031.42 Mb. Mean read length per dataset ranged from 1.01 to 11.12 kbp, with a mean of 4.43 kbp, median of 4.00 kbp and standard deviation of 1.87 kbp. 151 samples were monocultured isolates, while 50 contained multiple species and/or potential host contamination. The monocultured isolates were used for the majority of the tests here; the remaining 50 were used to test how the tools performed with more “difficult” samples.

27 bacterial species were included in the monoculture or metagenomic datasets, with *Escherichia coli* the most common species (40 samples) and *Staphylococcus* the most widely represented genus (40 samples across 7 species). On average, each species was represented by 6.3 samples (range=1-40 samples). For the purposes of statistical power, only antibiotic classes recorded as resistant in ASTs from at least 5 (∼2%) of our samples were considered (**Supplementary Table S5**). Of these, beta-lactam resistance was most common (52.24% of samples). The mean number of samples in our dataset resistant to each antibiotic class was 35.7 (SD=30.11). See **Supplementary Table S4** for full species and AST details.

We also assessed the distribution of positive and negative AST calls. Overall, at the level of individual antibiotic there were 1,427 AST calls, 38% positive (resistant or intermediate) and 62% negative (susceptible), whilst at the level of antibiotic class there were 760 AST calls, 46% positive and 54% negative, suggesting acceptable overall balance. **Supplementary Tables S11 and S12** show the distributions across each antibiotic and class, with some antibiotics or classes more evenly distributed than others. Balanced accuracy was used instead of accuracy in subsequent analyses here, because it averages sensitivity and specificity and therefore reduces the influence of imbalance.

### Neither basecalling accuracy mode nor sequencing chemistry significantly affects overall AMR prediction accuracy

A subset of 64 monocultured samples were basecalled with Dorado in three different accuracy modes: fast, high accuracy (HAC) and super accuracy (SUP). Each set of reads was then processed through the rest of the assembly, analysis and statistics pipeline, and the results stratified by basecalling accuracy.

**Figure 1** shows that no significant differences were found between the accuracy modes across any of the performance metrics, although there is a slight trend towards increased sensitivity from fast to HAC to SUP. All samples in our dataset were therefore carried forward for further analysis, regardless of original basecalling accuracy.

**Figure 1.**
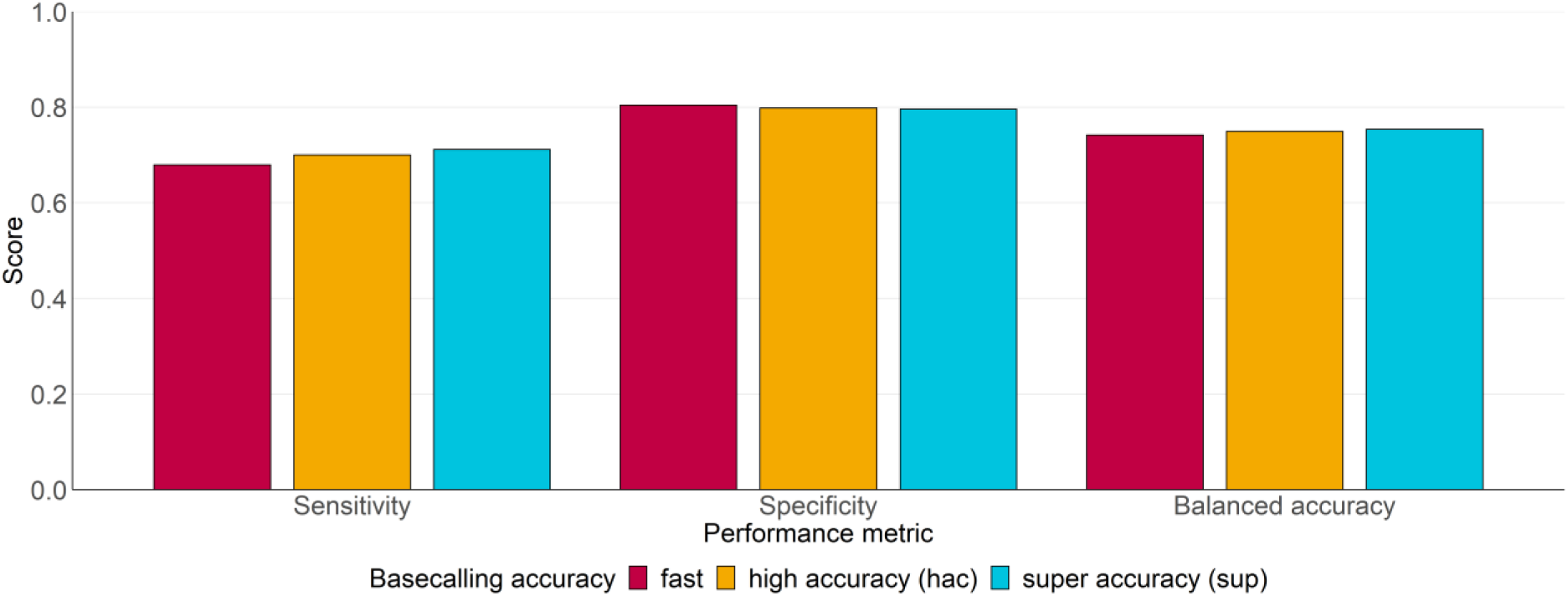
Performance metrics of our 64-sample subset which were basecalled with three different basecalling accuracy modes in Dorado. Metrics were calculated across all tools and all samples, then stratified by basecalling accuracy; fast, high accuracy (hac) and super accuracy (sup). No significant differences were observed, according to the Kruskal-Wallis test.

The results for a small subset of species-matched R10.4.1 and R9.4/R9.4.1 were compared to reveal any differences due to the use of older or newer sequencing chemistry or basecaller. No significant differences were seen in any performance metric between R10 and R9.

### Flye+Medaka produces the best balanced accuracy, whilst reads-only produce the best sensitivity across all tools

Performance metrics were calculated for all 151 monocultured samples across all tool/database combinations, then stratified by assembly strategy (“flye”, “medaka”, “miniasm” and “reads”). As shown in **Figure 2**, although no tool performed significantly better across all metrics, the reads-only strategy produced significantly better sensitivity than all other strategies, whilst the Flye assembly with or without Medaka polishing produced significantly better results than Miniasm for balanced accuracy, and significantly better specificity than the reads-only strategy. The Medaka and reads-only strategies were therefore taken forwards for more detailed analysis across all tools.

**Figure 2.**
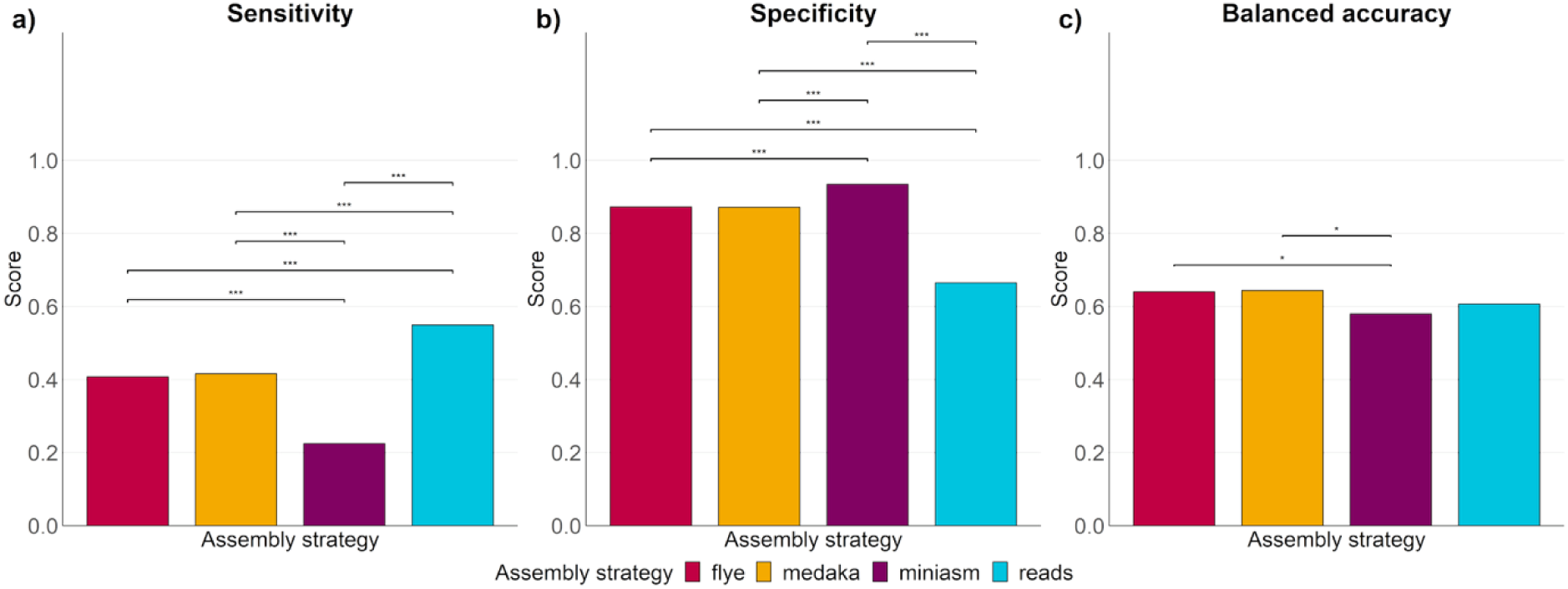
Overall performance metrics across 151 monoculture samples and all antimicrobial resistance detection tools, stratified by assembly strategy. a) Sensitivity, b) specificity, and c) balanced accuracy. “Flye” refers to assembly with Flye in nano-hq mode, “medaka” refers to assembly with Flye followed by polishing with Medaka, “miniasm” refers to rapid assembly with Miniasm, and “reads” refers to prediction directly from raw sequencing reads without an assembly step. Of the tools tested here, only AMR++, DeepARG and ResFinder include a read-only mode. Statistical significance was calculated using the Kruskal-Wallis test and Dunn’s post-hoc test; *p*-values indicate pairwise significance. *** indicates *p* < 0.0005, ** indicates *p* < 0.005, and * indicates *p* < 0.05.

### ResFinder-based tools produce the best overall balanced accuracy

The mean score for each performance metric across the 151 monocultured samples was calculated and stratified according to prediction strategy (tool, database, Medaka assembly vs reads) to determine the best performing tool overall. **Figure 3** shows the results for sensitivity, specificity and balanced accuracy.

**Figure 3.**
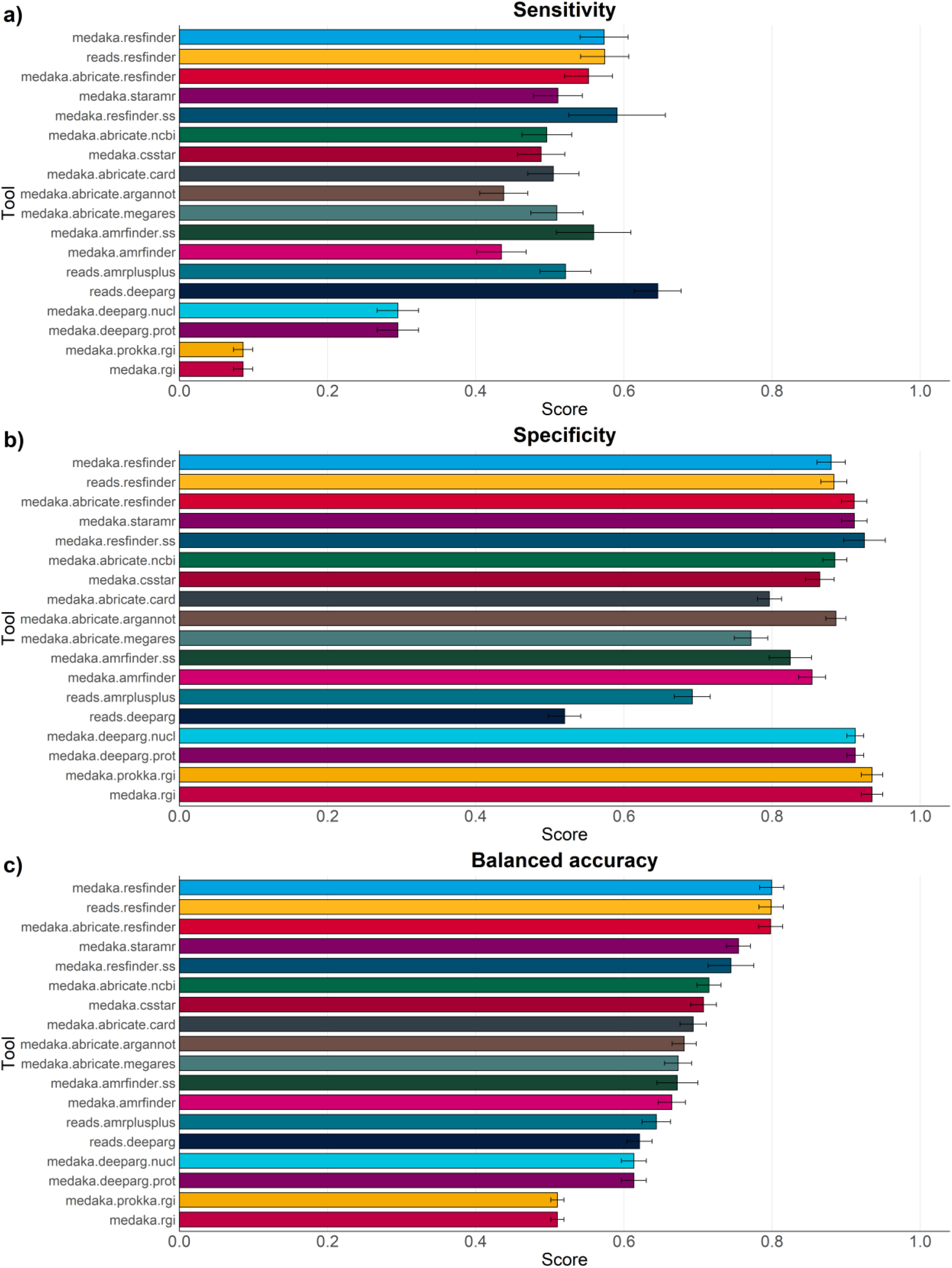
Performance metrics for each AMR prediction strategy, calculated at the level of specific antibiotics per sample then stratified by prediction strategy across the 151 monoculture samples. “Tool” names follow the general structure assembly/reads.tool.(database).(ss), where (database) is only relevant for those tools with multiple database options and (SS) is only relevant for those tools with a species-specific mode. “reads” indicates prediction directly from raw sequencing reads, whereas “medaka” indicates Flye assembly followed by Medaka polishing. a) shows the mean sensitivity, b) shows the mean specificity and c) shows the mean balanced accuracy. Error bars indicate standard error. Abbreviations: NCBI, National Center for Biotechnology Information; CARD, Comprehensive Antibiotic Resistance Database; ARG-ANNOT, Antibiotic Resistance Gene-ANNOTation; RGI, Resistance Gene Identifier; MEGARes, MEGARes antimicrobial resistance database; nucl, DeepARG’s nucleotide-only mode; prot, DeepARG’s annotation/protein mode.

For balanced accuracy, which considers both sensitivity and specificity, the top five strategies are all based on the ResFinder database, with the ResFinder tool itself and ABRicate with its own version of the ResFinder database performing better (0.80 ± 0.02 for ResFinder in read-based mode and both tools in assembly mode) than StarAMR (0.75 ± 0.02) with its own version of the ResFinder database.

Sensitivity scores were generally lower than specificity. The read-based DeepARG mode scored the highest at 0.65 ± 0.03. The lowest sensitivity scores were for both RGI strategies (assembly and annotations) at 0.09 ± 0.01, meaning that most AMR genes were not called.

Conversely, both RGI modes ranked the highest in specificity (0.93 ± 0.1 for assembly and annotation-based). However, as specificity relates to the absence of false positive calls, RGI scoring highly does not mean that it performed particularly well; rather, RGI made the fewest calls simply because it made so few positive calls (false or true) in general. Read-based DeepARG performed best in sensitivity, but worst in specificity (0.52 ± 0.02), indicating a high number of apparent false positives alongside the true positives. In contrast, the two modes of the ResFinder tool score relatively well for specificity: 0.88 ± 0.02 for both read-based and assembly.

### Using “species-specific” databases may increase the accuracy of AMR predictions

We investigated the species-specific mode of two of the AMR tools tested here: ResFinder and AMRFinderPlus, to test whether using databases of species-specific genes and point mutations improved the overall accuracy of AMR predictions. **Supplementary Table S4** shows the species for which species-specific modes were available for the two tools. For ResFinder, species-specific predictions were only available for three of the species in our dataset (n=56 monoculture samples), whereas for AMRFinderPlus they were available for an additional 11 (14 total species, n=122 monoculture samples). Compared to the non-species-specific results, the sensitivity and specificity of species-specific ResFinder improved slightly (0.57 to 0.59 and 0.88 to 0.92, respectively). For AMRFinderPlus, the species-specific sensitivity showed a larger increase (0.43 to 0.56), but the specificity decreased (0.85 to 0.82).

### Making predictions based on antibiotic classes improves sensitivity, but reduces specificity

We tested whether grouping antibiotics into their classes (**Supplementary Table S4**) produced better predictions, as some of the tools tested originally reported AMR on the level of class only. **Figure 4** shows the per-class results for balanced accuracy, sensitivity and specificity, stratified by tool/database/reads vs assembly. The strategies which performed best on the level of specific antibiotic also performed best here: ResFinder-based tools were the top five scorers in balanced accuracy, whilst read-based DeepARG performed best in sensitivity but worst in specificity. Compared to predictions for specific antibiotics, read-based and assembly-based ResFinder scored worse for specificity: 0.88 per specific antibiotic and 0.81 per antibiotic class. In contrast, sensitivity was significantly improved: 0.57 per specific antibiotic and 0.73 per antibiotic class for ResFinder, and 0.65 per antibiotic and 0.80 per class for DeepARG.

**Figure 4.**
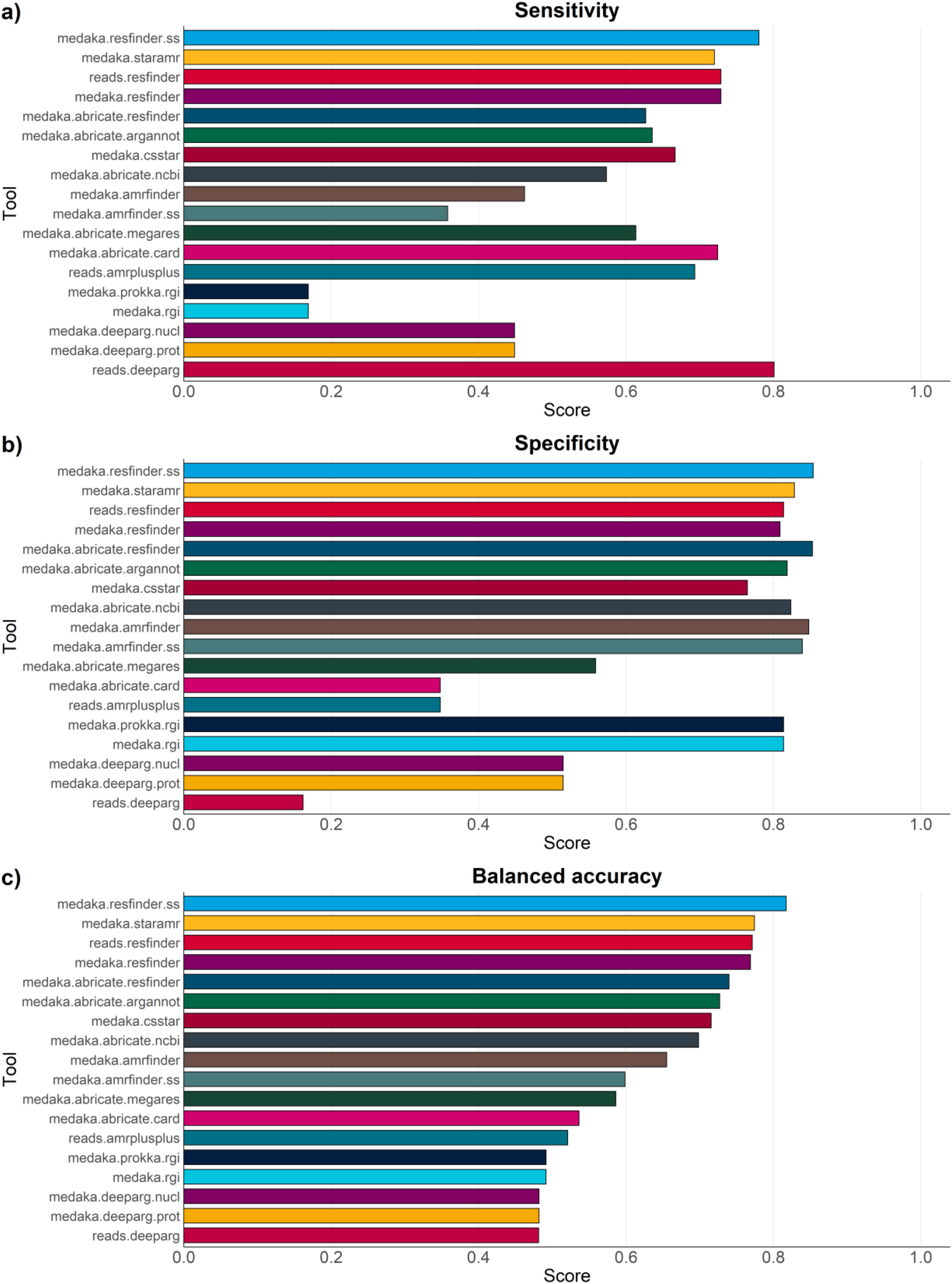
Performance metrics for each AMR prediction strategy, calculated at the level of antibiotic class and stratified by prediction strategy across the 151 monoculture samples. “Tool” names follow the general structure assembly/reads.tool.(database).(ss), where (database) is only relevant for those tools with multiple database options and (SS) is only relevant for those tools with a species-specific mode. “reads” indicates prediction directly from raw sequencing reads, whereas “medaka” indicates Flye assembly followed by Medaka polishing. a) shows the mean sensitivity, b) shows the mean specificity and c) shows the mean balanced accuracy. Error bars indicate Standard Error. Abbreviations: NCBI, National Center for Biotechnology Information; CARD, Comprehensive Antibiotic Resistance Database; ARG-ANNOT, Antibiotic Resistance Gene-ANNOTation; RGI, Resistance Gene Identifier; MEGARes, MEGARes antimicrobial resistance database; nucl, DeepARG’s nucleotide-only mode; prot, DeepARG’s annotation/protein mode.

### AMR predictions are more accurate for some antibiotic classes than for others

We investigated whether AMR predictions for some classes were more robust than for others. **Figure 5** shows extensive significant differences in sensitivity, specificity and balanced accuracy across the different classes. Of note for sensitivity are polypeptides and fluoroquinolones, which performed particularly poorly (0.07 and 0.22 respectively), and aminoglycosides and tetracycline, which performed the best (0.69 and 0.72 respectively). Glycopeptides, although tested in only a small number of samples, performed well for specificity (0.93) but more poorly for sensitivity (0.52).

**Figure 5.**
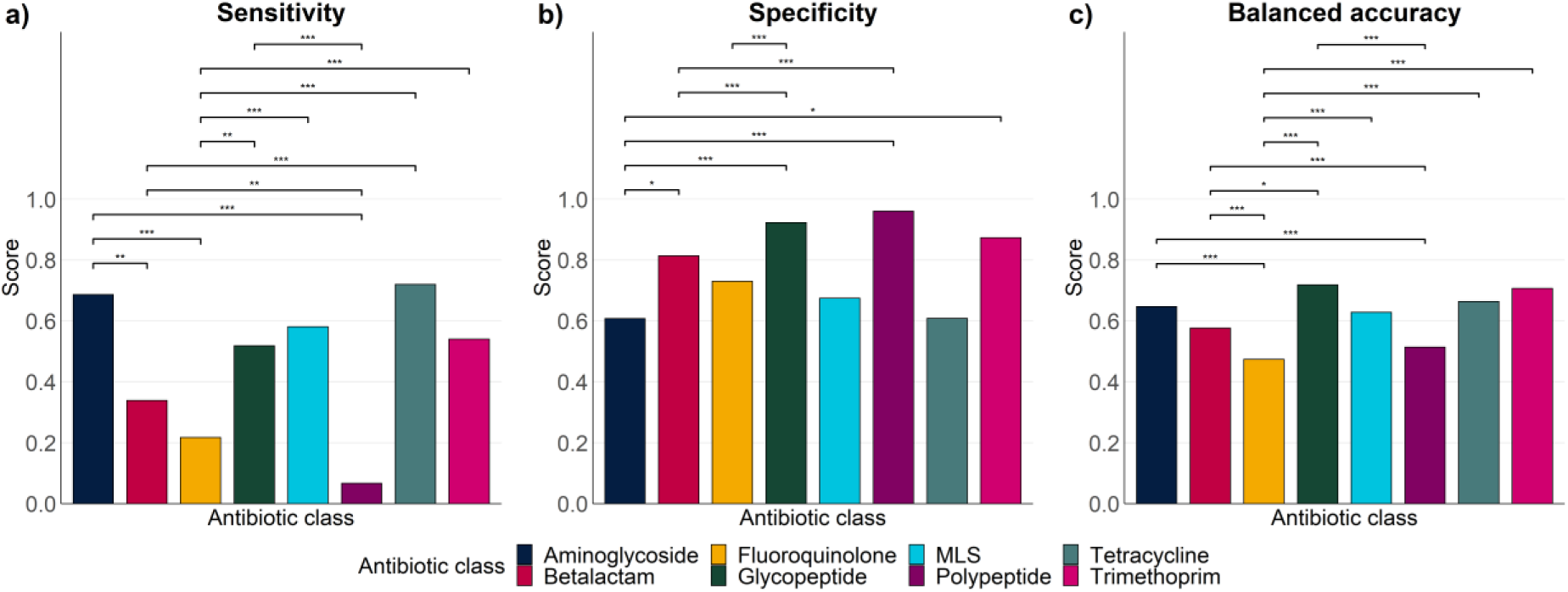
Overall performance metrics across 151 monoculture samples across all prediction strategies, stratified by antibiotic class. a) Mean sensitivity, b) mean specificity, and c) mean balanced accuracy. MLS indicates macrolide-lincosamide-streptogramin. Statistical significance was calculated using the Kruskal-Wallis test and Dunn’s post-hoc test; *p*-values indicate pairwise significance. *** indicates *p* < 0.0005, ** indicates *p* < 0.005, and * indicates *p* < 0.05.

### AMR predictions are more accurate for some species than for others

To further investigate the factors affecting the accuracy of our predictions, we calculated the mean sensitivity, specificity and balanced accuracy across the 151 monocultured samples for the overall best performing prediction strategy (assembly with Flye, polishing with Medaka, prediction with ResFinder), stratified by the genus of the original sample. **Figure 6** shows that the predictions for some species are significantly more accurate than for others. *Klebsiella* (n=5) scored 100% for sensitivity, meaning every instance of AMR was predicted correctly, while *Staphylococcus* (n=36) also scored well, with almost 90% sensitivity. Meanwhile, both *Pseudomonas* (n=18), *Haemophilus* (n=8) and *Mycobacterium* (n=10) scored very poorly for sensitivity; every instance of AMR in *Mycobacterium* was missed. Most genera performed better for specificity than for sensitivity, with *Mannheimia* (n=15), *Enterococcus* (n=11), *Mycobacterium* (n=10) and *Neisseria* (n=7) all scoring 100%, and *Escherichia* (n=44) and *Staphylococcus* (n=36) both scoring over 90%.

**Figure 6.**
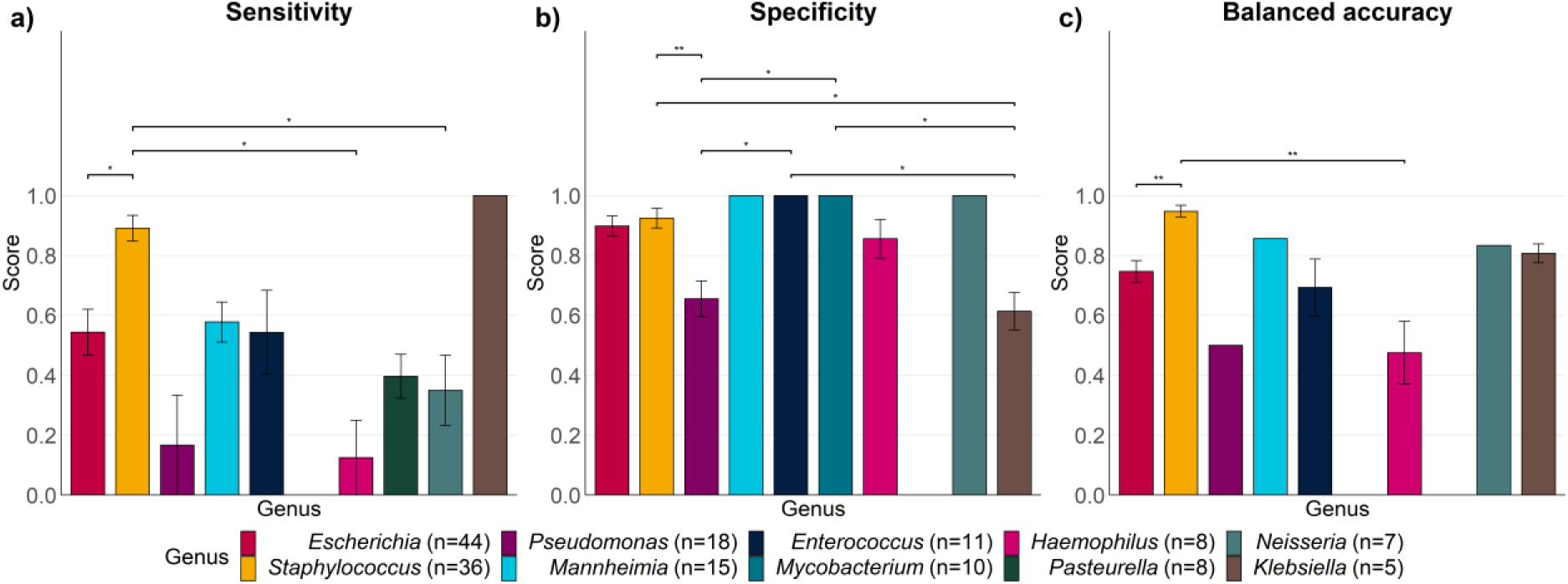
Overall performance metrics across 151 monoculture samples assembled with Flye, polished with Medaka, and screened for antimicrobial resistance using ResFinder, stratified by genus identified in the original sample. a) Mean sensitivity, b) mean specificity, and c) mean balanced accuracy. Where a genus scored 0 in either sensitivity or specificity, balanced accuracy could not be calculated. Error bars show standard error. The legend indicates the number of samples included for each genus. Statistical significance was calculated using the Kruskal-Wallis test and Dunn’s post-hoc test; *p*-values indicate pairwise significance. *** indicates *p* < 0.0005, ** indicates *p* < 0.005, and * indicates *p* < 0.05.

### AMR prediction tools for samples with mixed populations or host contamination perform similarly to more simple samples

To test how confounding factors such as host DNA contamination affected the prediction accuracy of our AMR tools, we processed 50 metagenomic/mixed species samples through the same analysis pipeline. This showed the same pattern as the monoculture results: ResFinder-based tools performed best for overall balanced accuracy whilst DeepARG performed best for sensitivity but worst for specificity. However, although the performance pattern remained the same the actual scores observed for the “difficult” samples were lower than those of the monocultured samples. Read-based ResFinder scored 0.75 ± 0.02 for balanced accuracy compared to 0.80 ± 0.02 for the monocultures, whilst DeepARG scored 0.58 ± 0.03 for sensitivity compared to 0.65 ± 0.03 for monocultures. Full results are available in the **Supplementary Materials** (**Supplementary Figure SF1**).

### Increasing starting data volume raises the overall accuracy of AMR predictions

The effect of data volume on the AMR prediction performance was assessed, by analysing a subset of our samples filtered to 500 Mb, 250 Mb, 100 Mb, 50 Mb and 25 Mb. **Figure 7** shows that a significant improvement is seen in the sensitivity and balanced accuracy of the AMR predictions from 25 Mb to 500 Mb. There is an upwards trend across all volumes in both sensitivity and balanced accuracy, although for both, the only statistically significant differences are between 25 Mb and each of the other volumes. The incremental improvements in sensitivity get smaller with additional data: the difference between 25 Mb and 50 Mb is 0.07 (0.38 to 0.45), compared to only 0.01 between 250 Mb and 500 Mb (0.51 to 0.52). All significant differences seen across specificity were again between 25 Mb and the other volumes (except 50 Mb); in this case, there is a significant decrease from 25 Mb to 500 Mb (0.88 to 0.82). This is likely due to overall fewer (true or negative) positive calls being made at lower data volumes due to poorer assembly quality. At higher data volumes, more positive calls are made, resulting in more true positives, but also more false positives.

**Figure 7.**
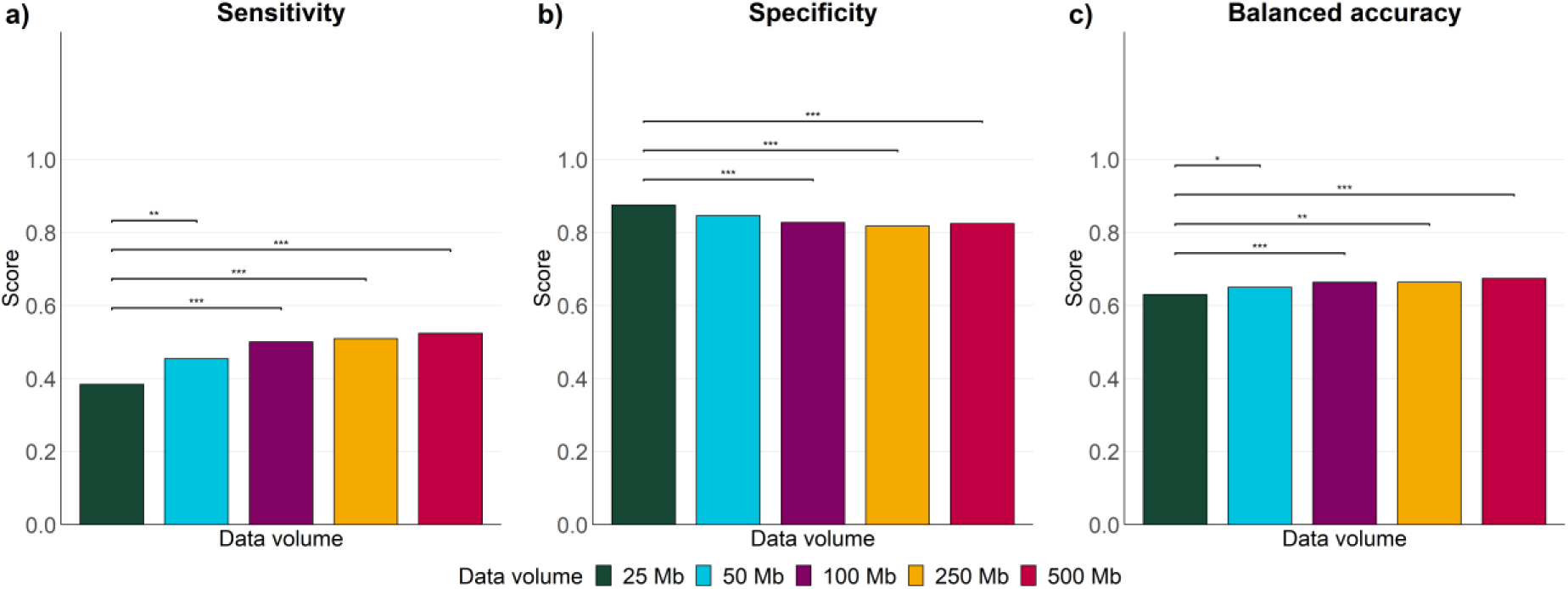
Performance metrics for the 61-sample subset filtered to different volumes of raw sequencing data using Filtlong, from 500 Mb down to 25 Mb. a) Sensitivity, b) specificity, and c) balanced accuracy. Metrics were calculated across all tools and samples, then stratified by starting data volume. Statistical significance was calculated using the Kruskal-Wallis test and Dunn’s post-hoc test; p-values indicate pairwise significance. For balanced accuracy, the Kruskal-Wallis test indicated a significant trend across all volumes, but no pairwise comparison was significant by Dunn’s post-hoc test. *** indicates p < 0.0005, ** indicates p < 0.005, and * indicates p < 0.05.

## 8. Discussion

### Benchmarking AMR prediction tool performance with ONT data could facilitate future clinical implementation

In this study, we set out to determine how well existing computational tools can detect AMR in genomic DNA datasets derived from ONT sequencers, by comparing *in silico* AMR predictions (WGS-based antimicrobial susceptibility predictions, WGS-ASP) with paired *in vitro* AST results. Although there are many *in silico* AMR detection tools available, the majority were originally designed with Illumina short reads in mind. Indeed, some commonly-used tools (e.g. ARIBA (84), SRST2 (85), Groot (86)) could not be tested here, as they are for short read data only. The new Amira tool is designed specifically for long reads; however, at the time of testing it was only available for a limited number of species and thus we did not include it (87).

The tools which can be used with ONT data were still therefore originally optimised to detect AMR genes and mutations based on the characteristics of Illumina reads. These are short (<500 bp) reads with high (often >99.5%) raw accuracy, and any errors tending to occur as substitutions at the end of reads (88). These tools therefore may not perform as well with reads from ONT sequencers, which are long (typically >1,000 bp) and historically had a higher raw error (1-5%), with errors tending to be insertions or deletions (indels) (89). Raw accuracy is much higher using the latest ONT sequencing chemistry and basecaller (R10.4.1 flow cells and Dorado commonly produce raw error <1% (90)). However, our comparison of a small subset of species-matched samples sequenced on R10.4.1 and R9.4/R9.4.1 showed no significant difference between the overall accuracy of AMR predictions from the latest chemistry and basecaller and those of their predecessors. Tools designed with an Illumina error profile in mind may therefore have still resulted in missed AMR calls. For example, the default mode of RGI only calls exact matches as hits, thus explaining its surprisingly poor performance here; even minimal indels could cause frameshifts in the gene sequences, which would prevent their accurate translation into protein sequences leading to false negative results (33). Applying RGI’s “--include_loose” flag could allow for higher sensitivity with ONT data.

With metagenomic ONT sequencing likely to used diagnostically in clinical settings, it will be vital that robust computational methods are used to identify species and detect AMR profiles. Accurately classifying AMR in genomic DNA datasets is also useful in other contexts, such as monitoring the environmental load of AMR-carrying bacteria within healthcare or One Health settings (91–94). Whilst there are well-established and maintained guidelines for the interpretation of *in vitro* lab-based ASTs (95), no such guidelines currently exist for interpreting WGS-ASP. Understanding the strengths and weaknesses of the possible strategies to detect AMR from genomic data will therefore be important when determining which methods to implement in clinical settings. Whilst benchmarking has previously been done for short reads (27) but there is less evidence for the accuracy of prediction strategies from ONT long read sequencing data, although we note that a very recent publication from Govender *et al.* benchmarked several of the same tools tested here with metagenomic samples from blood cultures, and also found that ResFinder produced the results most closely matched to AST (96). In addition, we here address the impact several important data steps prior to AMR prediction: namely, basecalling accuracy, volume of data, and assembly strategy.

It should be acknowledged that the majority of our comparisons here were carried out using monocultured isolate samples (n=151); there were simply far more datasets for this kind of sample available. Although our separate analysis of the metagenomic samples (n=50) showed slight decreases in the performance metrics compared to the monoculture samples (for example, the decrease of ResFinder’s balanced accuracy from 0.80 to 0.75), we observed the same overall patterns in tool performance. The results of the monoculture analysis can therefore be regarded as a cautious proxy for more complex samples. In addition, both datasets also included uneven contributions from multiple source studies, which differed in sample number, sequencing chemistry, basecaller version and accuracy mode, genera and AST panels. Source study composition may therefore have influenced our performance estimates and tool rankings. We stratified the results shown in **Figure 3** and **Supplementary Figure SF1** by original data source, and report the corresponding sample counts, TP/FP/TN/FN counts, sensitivity, specificity, and balanced accuracy in **Supplementary Table S13**. Overall, the source-stratified analysis supported the main interpretation of the pooled analysis: performance varied by tool/database and reflected sensitivity/specificity trade-offs.

### Choice of basecalling mode does not significantly affect AMR prediction accuracy, but choice of assembly strategy does

Before comparing the performance of the AMR prediction tools and databases, we optimised the strategy for preparing sample data. The first step in analysing ONT data is basecalling, i.e translating the raw electrical signal (the “squiggle”) into a DNA sequence. Early basecallers such as Albacore, which was used to basecall 34 of the samples in our dataset, did not have different basecalling modes. However, Guppy and Dorado have three accuracy modes: fast, high accuracy (HAC) and super accuracy (SUP). SUP basecalling requires much higher computing resources and takes an order of magnitude longer than fast basecalling, albeit with the benefit of significantly increased per-base accuracy in the resulting fastq reads. HAC basecalling is the middle-ground option between fast and SUP. ONT do not currently publish an expected read accuracy for fast basecalling, but quote a median raw read accuracy of 99.4% and 99.7% respectively for the latest HAC and SUP models (97).

SUP basecalling within a rapid diagnostic timeframe would require a GPU-powered computer or GridION, which may not be economically viable in all clinical settings. However, despite a slight trend in our results towards more accurate WGS-ASP from fast to HAC to SUP, the differences were not significant. Our results therefore indicate that even fast basecalling, which is feasible using a standard laptop with 2-4 CPUs and 8-16 Gb RAM, may produce AMR predictions with comparable accuracy to HAC or SUP basecalling. We note that our basecalling analysis was based on monocultured isolates; it is possible, therefore, that different basecalling accuracy modes may produce more significantly different results when applied to metagenomic samples.

The next step in data preparation after basecalling is assembly into contigs. A previous comparison of AMR prediction tools focussing on short read data included tools that analysed reads but not assembled contigs (27). Although assembling reads into contigs requires additional processing steps and therefore additional time, the processing of individual reads into a consensus sequence greatly improves the per-base accuracy (98). For ONT data, which may be less accurate than Illumina data at the read level, computing a consensus may therefore result in more accurate AMR predictions. Our results show that Miniasm, which exchanges accuracy for speed, performed the worst overall in sensitivity and balanced accuracy. Flye and Flye+Medaka produced results that were not significantly different from each other; this finding is unsurprising, since newer sequencing chemistries and basecallers may not need post-assembly polishing (99). However, since **Figure 2** does show a marginal improvement from Flye to Flye+Medaka, we chose to use the Medaka assemblies instead of Flye when analysing the performance of each AMR prediction tool separately.

### ResFinder DB produces results which most closely match AST, whilst DeepARG produces the highest sensitivity

We tested a variety of tools with a variety of databases, in both assembly and read-based modes. Where a tool allowed either assembly or read mode to be used, we trialled both. Likewise, where a tool allowed the selection of AMR databases (e.g. ABRicate), we trialled all available. ResFinder, CARD, MEGARes and NCBI’s AMRFinderPlus/NDARO are among the most commonly used AMR gene databases (100), and all were trialled with more than one tool. Our results show that, in terms of balanced accuracy between sensitivity and specificity, the ResFinder database produces WGS-ASP which most closely match the AST results, independent of the tool used to screen the database. Of the ResFinder-based tools, the standalone ResFinder tool performed best.

ResFinder performed almost exactly the same when used with assembled contigs or with raw reads, suggesting that important AMR information from the reads is not lost during consensus building. ResFinder performed best in species-specific mode, although the ResFinder command line tool only includes species-specific screening for a very limited number of species; here, we could only test it with *Escherichia coli*, *Enterococcus faecalis* and *Enterococcus faecium* (n=56). Notably, the ResFinder webserver (101) offers species-specific screening with PointFinder for far more species than the command line tool. In addition, the webserver is user-friendly and outputs an easily interpreted results report. One AMR prediction strategy for rapid diagnostics would therefore be to simply use the ResFinder webserver with either assembled fasta or raw fastq reads, selecting any species known to be in the sample. However, this relies on a shared public web service; analysis turnaround time may therefore vary with server load and concurrent usage. In addition, the need to upload genetic data to an online tool could raise ethical concerns, particularly when working with metagenomic samples originally from humans, which may still contain human DNA sequences.

Meanwhile, DeepARG with raw reads displayed the highest sensitivity here, albeit with the lowest specificity. Although the results of DeepARG are not the closest match to the AST results, therefore, DeepARG detects the overall highest number of AMR genes, and may represent a strong choice for surveillance of the full acquired resistance gene load in metagenomic samples.

### How accurate are the results of “gold standard” tests?

AST remains the most appropriate reference standard for benchmarking *in silico* AMR prediction, because it directly measures the resistance phenotype *in vitro*. However, phenotypic susceptibility testing is not error-free, with reproducibility and other outcomes ranging from 90-99% (102). Phenotypic AST results can vary with method, inoculum, media, incubation conditions, endpoint reading, and breakpoint interpretation, thus single AST results should not be interpreted as a fixed “ground truth”, a point which was emphasised in the recent European Committee on Antimicrobial Susceptibility Testing (EUCAST) update on WGS-ASP (103, 104). Variability close to clinical breakpoints is particularly important, and small technical or biological differences can alter whether an isolate is categorised as susceptible, intermediate, or resistant.

Consequently, although false positive or false negative results may reflect real errors by the prediction tool, some apparent false positive or false negative *in silico* predictions may instead occur due to limitations in the AST methods used, with limited ability to distinguish between the two. Isolates with known resistance determinants can therefore be deemed susceptible by ASTs, because a gene is present, but not phenotypically expressed at levels deemed “resistant” under laboratory conditions (105, 106). Such “silent” or poorly expressed resistance genes still reflect a latent AMR potential that could be clinically relevant, particularly if expression is induced *in vivo* or under antibiotic selection pressure.

These sources of discordance may affect the apparent performance of all tools evaluated here, particularly because our datasets were gathered from many different original studies which may have applied different AST methods in different laboratory settings. In particular, tools or databases that make broader genotype-to-phenotype assignments may generate more apparent false positives relative to AST. If these “false positive” calls are actually due to variability in the way AST breakpoints were interpreted, or under-expression of genes in laboratory conditions, a tool with apparently poor specificity may therefore actually represent a more conservative option for complementing clinical decisions.

Handling of “intermediate” resistance calls can also introduce apparent genotype-phenotype discordance. The EUCAST guidelines no longer group intermediate calls with resistant calls; instead, intermediate calls are treated as “susceptible, but requires longer exposure”, although it has been argued that this increases the rates of both false positive and false negative AST calls (107, 108). Here, we grouped resistant and intermediate phenotypes for the purposes of benchmarking genotype-based detection of reduced susceptibility, treating only “susceptible” phenotypes as truly susceptible. This means that if a tool detected no AMR determinants in an “intermediate” sample, we would deem this a false negative, but under the EUCAST guidelines it would be called susceptible, and therefore a true negative. Nonetheless, as only 20 of the 1,427 AST calls analysed here were “intermediate”, the classification of these as either resistant or susceptible likely had minimal impact on our overall results.

Therefore, although AST was retained as the “ground truth” for this benchmarking study, performance metrics should be interpreted as agreement with the available phenotypic test results, rather than as an absolute measure of biological truth.

### Is *in silico* AMR prediction ready for use in a clinical setting with ONT data?

In this study, we identified an optimal strategy for preparing ONT data for AMR prediction, and compared the performance of AMR prediction tools and databases. Tools using the ResFinder database strike the best balance between sensitivity and specificity, while DeepARG performs the best in terms of just sensitivity. However, even the best performing tool missed some AMR reported by phenotypic AST.

Although no diagnostic test is perfect, in order for a test to be trusted for making clinical decisions, its accuracy would ideally approach 100% (109). DeepARG, the best tool for sensitivity, detected only 65% of the AMR in our AST results. A number of factors may have led to this. Our dataset represented a wide variety of sequencing qualities, with some samples dating back to 2019 or earlier, sequenced on less accurate flow cells and basecalled with less accurate algorithms. However, our results suggest that neither basecalling accuracy nor sequencing chemistry significantly affected the accuracy of WGS-ASP. Conversely, some of our samples had minimal data, and our data volume test shows that even 500 Mb might not capture all AMR present in a sample.

Most importantly, a variety of genera were included in our samples. We included these to ensure our testing was representative of the spectrum of pathogens that could be present in clinical mWGS samples. Our results showed that whilst some genera scored highly for sensitivity, specificity and balanced accuracy, others scored poorly, particularly for sensitivity. The genera which scored the worst for sensitivity included those known to perform poorly with standard acquired resistance gene databases, such as *Pseudomonas*, in which resistance is usually a result of chromosomal variants, efflux, porin changes, and regulatory mechanisms, and *Haemophilus*, which accumulates resistance almost entirely through single nucleotide polymorphisms and indels (110, 111). *Mycobacterium* in particular is known to acquire resistance using mechanisms not usually represented in standard gene databases, and prediction of AMR tends to require the use of *Mycobacterium*-specific tools (112, 113). We may conclude, therefore, that the overall performance of the tools analysed here, which largely did not include mutation-based detection, has been influenced by the inclusion of genera for which gene-based AMR detection is difficult; for those genera whose resistance is based mainly on acquired genes, such as *Klebsiella*, the predictions from the tools tested here are much more reliable.

Predicting AMR based on antibiotic class instead of specific antibiotic may lead to better sensitivity; our results showed an improvement from 65% to 80% for DeepARG. Using this strategy, if a gene was detected which was related to AMR in a specific antibiotic, all other antibiotics from that same class would automatically be excluded. For example, if a gene related to gentamicin resistance was detected, all aminoglycosides would be excluded and an entirely different antibiotic class would need to be used. However, predictive antibiotic susceptibility is variable between classes, bacterial species and AMR genes. In our study, performance for aminoglycosides, glycopeptides, tetracyclines and trimethoprim were reasonably good, whilst that for beta-lactams was less impressive, and performance for polypeptides and fluoroquinolones was very poor. In genomic terms, this may reflect both gaps in AMR databases as well as resistance based on mutations rather than acquired genes (as seen in fluoroquinolone resistance) (114). Most tools do not offer a species-specific mode or the option to screen for point mutations, and those that do (such as ResFinder) offer it only for a limited group of species. Consequently, sensitivity to AMR caused by point mutations will be low. Care should therefore be taken with any class-based strategy to avoid excluding antibiotics that may be effective, or including those where resistance is likely. This is particularly important in serious infections and/or multi-drug-resistant scenarios where clinical outcomes depend on a rapid and accurate choice of antibiotic, and fewer antibiotics are available.

### Conclusions

We have here detailed several important findings for choosing an optimal pipeline for AMR prediction from nanopore data: choice of basecaller accuracy mode seems to exert little influence on overall WGS-ASP accuracy; increasing data volume results in more accurate WGS-ASP but with diminishing returns, and a strategy using either raw reads or contigs assembled with Flye (with or without Medaka polishing) will produce WGS-ASP with similar accuracy. However, our results indicate that limitations remain for users considering the implementation of AMR prediction tools and databases with nanopore long reads in clinical settings. This finding is consistent with an earlier study with Illumina short reads, and the recommendations of the recent EUCAST WGS-ASP report (27, 104). Universal guidelines for how *in silico* predictions should be interpreted for each species, antibiotic and antibiotic class are needed. Species-specific tools and databases may be required, and indeed already exist for species like *M. tuberculosis* and, increasingly, *P. aeruginosa* (110, 112, 113, 115).

In the meantime, the results of *in silico* predictions should be used with caution, with clinical decisions being made on a case-by-case basis, taking into account the case history, pathogen species, antibiotic class and available drugs, and bearing in mind that prediction tools work better for some bacteria and some antibiotics than for others. For example, a prediction of vancomycin (glycopeptide) resistance in *Staphylococcus* could be regarded with strong confidence, whereas the absence of detection of fluoroquinolone resistance in *Pseudomonas* would not necessarily reliably suggest susceptibility. Overall, based on our findings, we suggest that *in silico* AMR prediction using mWGS is ready to support and augment the results of *in vitro* AST, particularly in rapid patient-side diagnosis and treatment decisions, but it is not yet ready to replace it.

## Supporting information

Supplementary figures, methods and discussion

## 9. Author statements

### 9.1 Author contributions

Conceptualization: N.R.; Data curation: N.R., A.S.L., G.K.P.; Formal analysis: N.R.; Funding acquisition: J.R.F.; Investigation: N.R., A.S.L., R.E., M.K.; Methodology: N.R.; Project administration: D.N.C, J.R.F.; Resources: N.R., A.S.L., G.K.P., T.N.; Supervision: D.G., D.N.C, J.R.F.; Validation: N.R.; Visualization: N.R.; Writing -original draft: N.R., D.N.C., J.R.F.; Writing -review & editing: N.R., A.S.L., R.E., M.K., G.K.P., T.N., D.G., D.N.C, J.R.F.

### 9.4 Ethical approval

No ethical approval was required for this study.

## 9.5 Acknowledgements

The authors are grateful to the Easter Bush Pathology microbiology laboratory for the storage and provision of the samples sequenced specifically for this study. In addition, we thank the authors of the original studies for making their data available for the remainder of the samples analysed here.

