## Supplementary figures, methods and discussion for "Benchmarking nanopore-based strategies for antimicrobial resistance prediction"

#### Supplementary Figures

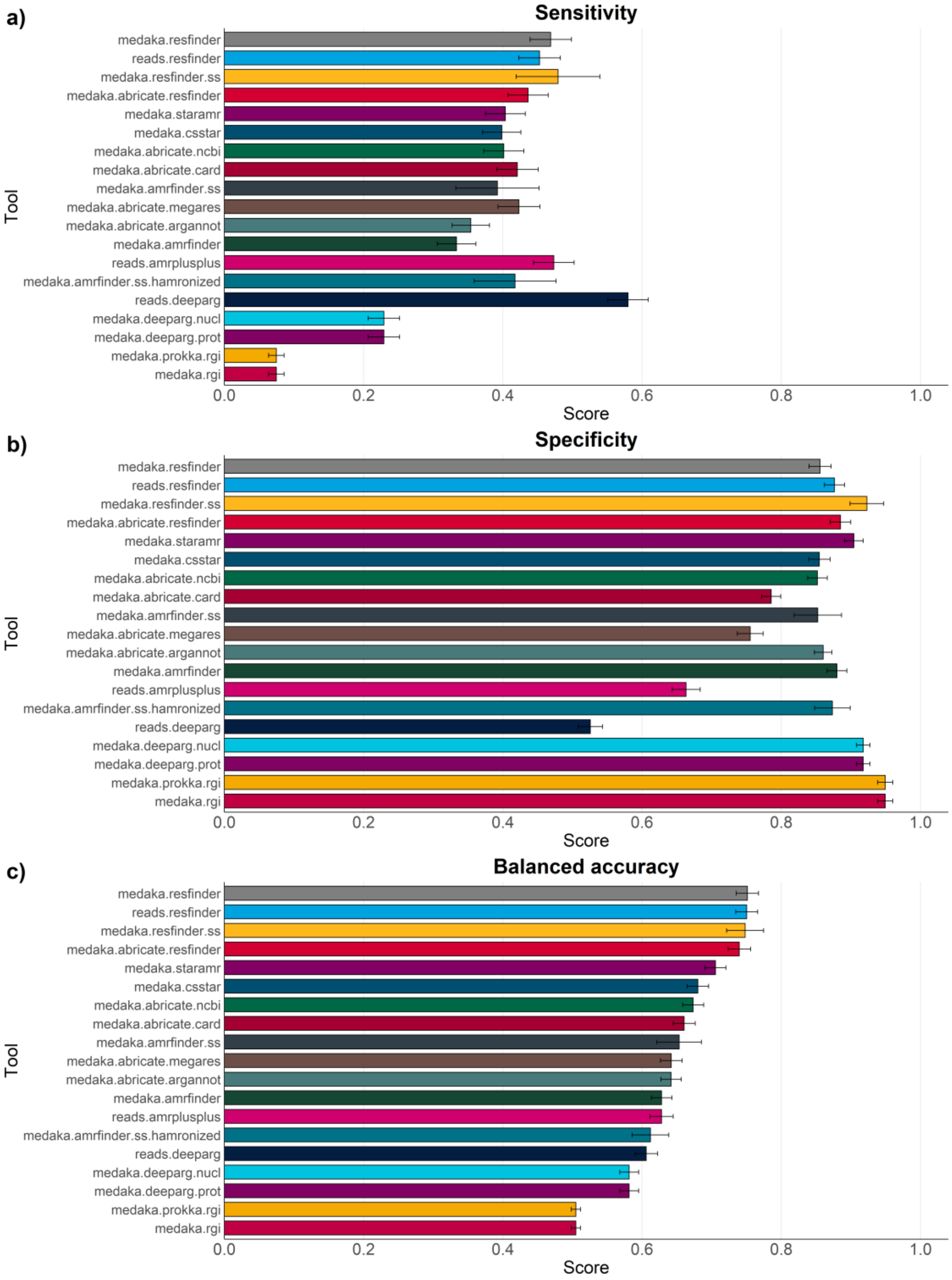


**Figure SF1. Performance metrics for each AMR prediction strategy, calculated at the level of individual antibiotic and stratified by prediction strategy across the 50 "difficult" metagenomic samples**. "Tool" names follow the general structure assembly/reads.tool.(database).(ss), where (database) is only relevant for those tools with multiple database options and (SS) is only relevant for those tools with a species-specific mode. "reads" indicates prediction directly from raw sequencing reads, whereas "medaka" indicates Flye assembly followed by Medaka polishing. a) shows the mean sensitivity, b) shows the mean specificity and c) shows the mean balanced accuracy. Error bars indicate Standard Error.

Abbreviations: NCBI, National Center for Biotechnology Information; CARD, Comprehensive Antibiotic Resistance Database; ARG-ANNOT, Antibiotic Resistance Gene-ANNOTation; RGI, Resistance Gene Identifier; MEGARes, MEGARes antimicrobial resistance database; nucl, DeepARG's nucleotide-only mode; prot, DeepARG's annotation/protein mode.

#### Supplementary Methods

##### SM1. DNA extraction, library preparation and sequencing

A set of 64 additional isolates (**Supplementary Table S7**) were sequenced specifically for this study, representing species not otherwise found in the literature search described above. Bacteria were grown overnight on tryptic soy agar (TSA, Oxoid ThermoFisher, Massachusetts, USA) plates at 37 °C, then a single colony was transferred into tryptic soy broth (TSB, Oxoid) and incubated overnight at 37 °C with shaking. 2.5 ml of overnight culture was pelleted by centrifuging for 3 min at 16 000 g, then resuspended in 160 µl 50 mM Tris, 10 mM EDTA, pH 8.0. 20 µl of metapolyzyme (3.3 mg $ml^{-1}$) was added, and the solution was mixed by flicking. The solution was then incubated on a thermomixer for 1 h at 37 °C with 900 r.p.m. shaking to lyse the bacterial cell walls. Following cell wall lysis, DNA was purified by following the MagAttract HMW DNA (Qiagen, Hilden, Germany) Gram-positive protocol as per the manufacturer's instructions, eluting into 50 µl nuclease-free water and incubating for 3 minutes at 50°C with 1,400 RPM shaking.

The DNA was cleaned up using the ProNex Size-Selective Purification System (Promega, Wisconsin, USA) according to manufacturer's instructions with the following modifications: we used a starting volume 50 µl DNA and 200 µl ProNex beads (1:4 ratio), and eluted into 20 µl nuclease-free water which was incubated at 50°C with 400 rpm mixing on a thermomixer for 10 minutes.

DNA concentrations were quantified using the Qubit dsDNA HS kit according to the manufacturer's instructions. Where necessary, the samples were diluted with nuclease-free water to achieve a mass of 200-250 ng in 10 µl, then sequencing libraries were prepared using the Rapid Barcoding kit (Oxford Nanopore Technologies, Oxford, UK), SQK-RBK004 or SQK-114.24. Samples were then sequenced on R9.4.1 (for SQK-RBK004) or R10.4.1 (for SQK-RBK114.24) flow cells for up to 72h on a GridION device with real-time super accurate basecalling and barcoding (see **Supplementary Table S1** for details of which samples were processed with which kit and which basecaller version was used for each flow cell). The read sets for these isolates were then added to those collated from our literature search, and all data was processed using the following preparation and analysis pipeline.

##### SM2. Assembly and polishing tools tested here

**Miniasm** is one of the fastest assembly tools available, and uses a simple all-vs-all alignment step of the reads to build a consensus sequence, without any further processing (1). **Flye** is a widely used more thorough assembly tool based on a multi-step repeat graph-based consensus building method (2). **Medaka** is an ONT polishing tool which creates a pile-up of the raw reads to a draft consensus (e.g. from Flye), then uses a neural network-based algorithm to output an improved consensus sequence (3).

##### SM3. AMR tools tested here

**ABRicate** (4) is an alignment-based tool which uses BLAST (5) to screen assembled contigs for the presence of AMR genes in a user-specified database. Databases available include **NCBI AMRFinderPlus** (here referred to as "NCBI"), **CARD**, **ARG-ANNOT**, **ResFinder** and **MEGARes**. The versions of the databases used here (from January 2025) included 5,386, 2,631, 2,223, 3,077 and 6,635 AMR gene sequences, respectively. ABRicate is a widely used tool, but does not screen for point mutations, and does not have a "read only" mode. The output of ABRicate is a .tsv. The details contained in the .tsv vary according to the database used. For example, ABRicate with ResFinder outputs the name and symbol of the genes detected, along with the predicted antibiotic resistance phenotypes, whereas ABRicate with MEGARes outputs only the MEGARes accessions of the genes detected.

**AMRFinderPlus** (6) is an alignment-based tool which can use either BLAST or HMMs to screen assembled contigs, annotated proteins or sequencing reads for the presence of AMR genes in the **NCBI AMRFinderPlus** database. The version of the AMRFinderPlus database used here (2025-03-25.1) included 7,586 AMR proteins and 1,429 point mutations. To screen for point mutations, the species-specific mode must be used. The output of AMRFinderPlus is a .tsv, which contains the name and symbol of the genes detected, along with the predicted antibiotic resistance phenotypes.

**ResFinder** (7) is an alignment-based tool which uses BLAST and the *k*-mer-based KMA (8) to screen assembled contigs or sequencing reads for the presence of AMR genes in the **ResFinder** database. The version of the ResFinder database used here (2.6.0) included 3,213 sequences. ResFinder also includes a species-specific mode which uses the **PointFinder** database alongside the ResFinder database to include species-specific AMR-related point mutations in a limited number of species. The output of ResFinder is a .txt, which contains the name and symbol of the genes detected, along with the predicted antibiotic resistance phenotypes.

**c-SSTAR** (9) is the command line interface version of SSTAR, an alignment-based tool which uses BLAST to scan for the presence of AMR genes in assembled contigs, using the **ResGANNOT** database. The ResGANNOT database is a custom combination of the ResFinder and ARG-ANNOT databases. The ResGANNOT database used here included 1,007 sequences. The output of c-SSTAR is a .tsv, which contains the name and symbol of the genes detected.

**RGI** (Resistance Gene Identifier) (10) is an alignment-based tool which uses BLAST to screen predicted protein sequences for AMR variants, using the **CARD** database. The version of the CARD database used here (4.0.1) included 6,442 reference sequences. RGI accepts two inputs: assembled contigs or annotated protein sequences. If assembled contigs are submitted, Prodigal (11) is first used to predict open reading frames (ORFs) which are then scanned with BLAST. The output of RGI is a .txt, which contains the name and symbol of the genes detected, along with the predicted antibiotic resistance phenotypes.

**StarAMR** (12) is another alignment-based tool which uses BLAST to screen assembled contigs for AMR genes using the ResFinder database. The version of the ResFinder database used by StarAMR here (from August 2024) included 3,200 sequences. The output of StarAMR is a .tsv, which contains the name and symbol of the genes detected, along with the predicted antibiotic resistance phenotypes.

**DeepARG** (13) takes a deep learning approach combined with alignment with Diamond (14) to scan for AMR variants in either "LS" (long sequence) or "SS" (short sequence) mode. Each mode has its own deep learning model, which scan the DeepARG-DB, a combination of the CARD, ARDB (15) and UNIPROT (16) database built during the development of the models. Here, "LS" mode was used to scan for AMR in both nucleotide and amino acid annotations, whilst "SS" mode was used to scan for AMR in our raw fastq reads. The output of DeepARG is a .tsv, which contains the name and symbol of the genes detected, along with the predicted class of resistance phenotypes (e.g. "beta-lactam" or "multidrug").

**AMR++** (17) is a Nextflow pipeline-based tool built to use with raw read sets and the MEGARes database. Here we employed just the "resistome" pipeline, which uses BWA (18) to align reads to the database. The version of MEGARes used by AMR++ here (3.0) included 8,733 sequences. The output of AMR++ is a .tsv, which contains the MEGARes accession of the genes detected.

**SM4. Data reconfiguration and standardisation for performance metric calculation**

Each tool outputs its results in a different format (for example, .json, .txt). In addition, the level of detail reported varies between tools, with some tools listing only gene names, others listing a gene name and a related antibiotic class, and others listing gene name, antibiotic class and the specific antibiotics predicted to be resistant as a result of the gene (for more detail, see **Supplementary Material SM3**). Some data reconfiguration was therefore required to standardise the way results were reported.

HAMRonization v1.1.9 (53) was used to convert the results of all tools into a consistent tabular form**.** Next, two custom python scripts were used (both available from this manuscript's Github repository) for each HAMRonization output. gene_to_class.py attempts to match gene names between the HAMRonized results and the gene names in the NCBI's reference gene catalogue, reporting back the antibiotic class related to each gene in a new column, "antibiotic_class". gene_to_phenotype.py goes one step further and attempts to match the gene names from the HAMRonized results with gene names in the phenotypes.txt file from the ResFinder database, reporting back the specific antibiotics to which each gene is predicted to convey resistance in the "predicted_phenotype"column.

When the results were analysed to compare the tool outputs with our "ground truth" AST results (**Supplementary Table S4**), if a tool reported its own specific antibiotics or antibiotic class (in the "antimicrobial_agent" or "drug_class" columns of the HAMRonization output tables), these took precedence over the results of gene_to_class.py and/or gene_to_phenotype.py.

See **Supplementary Materials SD1** for a further discussion of data and nomenclature issues encountered.

### Supplementary Discussion

#### SD1. Inconsistencies in nomenclature and reporting hinder comparison efforts

Throughout this study, we encountered some issues which made comparisons either between different tools or between tools and our ground truth difficult. Firstly, when gathering our datasets, we discovered inconsistencies in the way AST results are presented and/or abbreviated. The former is more of an inconvenience than a true problem, as it means results have to be transcribed into the same formats in order to be compared. Inconsistencies in antibiotic abbreviations and nomenclature, however, is more than an inconvenience, as they could lead to incorrect results in our ground truth tables. One such example is the use of the abbreviation “FLU”, which could equally refer to either fluoroquinolone or flucloxacillin, which are completely unrelated but commonly used antibiotics in different antibiotic classes. Without knowing which antibiotic an abbreviation refers to, having an AST result is meaningless.

Another nomenclature issue is the existence of multiple names for the same antibiotic: Tazocin and Piperacilin/Tazobactam are the same drug, and the brand name and generic name are used with equal frequency. Likewise, Augmentin is co-amoxiclav, which is also amoxicillin+clavulanic acid. Issues can also arise with the cephalosporins and sulfonamides, which can be spelled with either “ph” or “f” (e.g. cefalotin vs cephalotin). Ontologies like ARO (the Antibiotic Resistance Ontology), part of the CARD database, aim to standardise the way in which AMR genes, mutations and phenotypes are named (10). However, no similar ontology seems to exist to document and standardise the naming of the antibiotics themselves. Likewise, no universal rules are used for how each antibiotic should be abbreviated when presenting AST results, although the European Committee on Antimicrobial Susceptibility Testing (EUCAST) published thorough guidelines in 2018 (19). Either way, good practice would suggest defining abbreviations when publishing results.

A similar issue is that different AMR prediction tools present their results in very different formats. Some output their results as a .json file, others as a.tsv or .txt file. hAMRonization (20) is a recent tool which aims to standardise the outputs of a variety of popular tools, and has gone some way towards alleviating this issue. However, some problems remain. The amount of detail given by each tool also varies greatly, partly dependent on which database they use. For example, MEGARes-based tools list only a shortened gene name, such as blaTEM (or even just TEM, for ABRicate with MEGARes) instead of blaTEM-4. Different blaTEM genes confer resistance to slightly different antibiotics; for example, blaTEM-2 is linked with resistance to Amoxicillin, Ampicillin, Cephalothin, Piperacillin and Ticarcillin, whilst blaTEM-4 is linked with resistance to Amoxicillin, Ampicillin, Aztreonam, Cefepime, Cefotaxime, Ceftazidime, Ceftriaxone, Piperacillin and Ticarcillin. Vagueness in gene names could therefore lead to too few or too many resistances being predicted, thereby resulting in either false positives or false negatives.
